# Aerobic GenX Defluorination by a Wastewater *Cladosporium halotolerans* Strain with a Genomically Expanded Haloacid Dehalogenase Repertoire

**DOI:** 10.64898/2026.07.29.741450

**Authors:** Esau De la Vega-Camarillo, Gayan Abeysinghe, Saurav Kumar Mathur, Rashmi Singla, Shravan Parunandi, Jorge Arreola-Vargas, Smriti Shankar, Sanjay Antony-Babu, Won Bo Shim

## Abstract

Per- and polyfluoroalkyl substances (PFAS) such as GenX (HFPO-DA) are aerobically recalcitrant contaminants for which biological treatment options remain scarce; the best-characterized microbial degraders require strictly anaerobic conditions and external cofactors. We isolated *Cladosporium halotolerans* strain CsHGX-1 from activated sludge at a municipal wastewater treatment plant using GenX as the sole carbon source. Whole-genome sequencing (32.4 Mb; 11,201 genes) revealed a PFAS-degradation gene repertoire substantially expanded relative to congeneric *Cladosporium* species, including 26 dehalogenases (four type-II haloacid dehalogenases, HADs), 141 cytochrome P450s, and 558 esterases/hydrolases. Under aerobic conditions with GenX (50 mg L-1) as the sole carbon source, strain CsHGX-1 removed 47.6 ± 1.7% of GenX within 48 h, accompanied by fluoride release (0.079 ± 0.018 mM) that was absent in abiotic controls, confirming genuine C-F bond cleavage. Time-resolved RNA sequencing (0, 6, 24, 48 h; n = 6 biological replicates) revealed a phase-structured transcriptional program: oxidative genes, including cytochrome P450s, peaked first (6 h; up to 25.4-fold), hydroxylation and reactive-oxygen-species-management genes peaked next (24 h; up to 33.5-fold), and the three type-II HAD genes peaked last (48 h; up to 50.9-fold), coincident with fluoride accumulation. A parallel resazurin metabolic assay over 5 days confirmed sustained catabolic activity in GenX-exposed cultures relative to controls (1.37-1.52-fold; p ≤ 0.003). These findings identify strain CsHGX-1 as, to our knowledge, the first Ascomycete fungus for which genomic and transcriptomic evidence links oxidative activation to haloacid-dehalogenase-mediated defluorination of an aerobically recalcitrant PFAS, extending the known diversity of fungal PFAS degraders beyond Basidiomycota white-rot taxa.

**IMPORTANCE:** GenX is a PFAS “replacement” chemical that the U.S. Environmental Protection Agency added to its list of hazardous constituents in 2024, yet no aerobic biological treatment exists for it: every well-characterized microbial degrader requires oxygen-free conditions and added cofactors. We show that a fungus recovered from ordinary wastewater sludge breaks down GenX while using oxygen, the same conditions already used in conventional treatment plants, with no nutrient or reductant supplementation. Genome sequencing showed why this strain is unusual: it carries far more dehalogenase and cytochrome P450 genes than its close relatives. Time-course RNA sequencing showed these genes switch on in a defined order, oxidation first, then carbon-fluorine bond cleavage, matching the appearance of free fluoride in the culture. This links genome content to a functional outcome in an Ascomycete fungus, suggesting aerobic fungal defluorinators may already be present, unrecognized, in engineered wastewater systems.

## OBSERVATION

### A wastewater fungus with a genomically expanded dehalogenase repertoire

Internal transcribed spacer (ITS) rDNA sequencing and maximum-likelihood phylogenetic reconstruction placed strain CsHGX-1 within the *Cladosporium halotolerans* species complex (99.2% identity to CBS 119424), as a distinct lineage sister to *C. sphaerospermum* (bootstrap 89; Figure 1a). This places the strain in Dothideomycetes, a class historically unexplored for halogen metabolism; nearly all previously characterized fungal PFAS degraders instead belong to Basidiomycota white-rot taxa such as *Phanerochaete chrysosporium* and *Trametes versicolor* (7, 8). Whole-genome sequencing (32.4 Mb; 94 contigs; N50 = 1.00 Mb; GC = 51.3%; 11,201 genes) revealed a PFAS-degradation gene repertoire markedly enriched relative to 12 other fungal genomes spanning Cladosporiaceae, other Ascomycota, and Basidiomycota white-rot taxa: 26 dehalogenases (1–3 in *Cladosporium* congeners), 141 cytochrome P450s (1.26% of the proteome, comparable to the 149 CYP450 genes of *P. chrysosporium* despite a smaller genome; 9), and 558 esterases/hydrolases (Figure 1b,c; Table S5). Among enzyme classes implicated in PFAS catabolism, only haloacid dehalogenases are known to catalyze direct hydrolytic C-F bond cleavage, making the four type-II HADs identified here the most functionally consequential feature of this genome (10).

**FIGURE 1.**
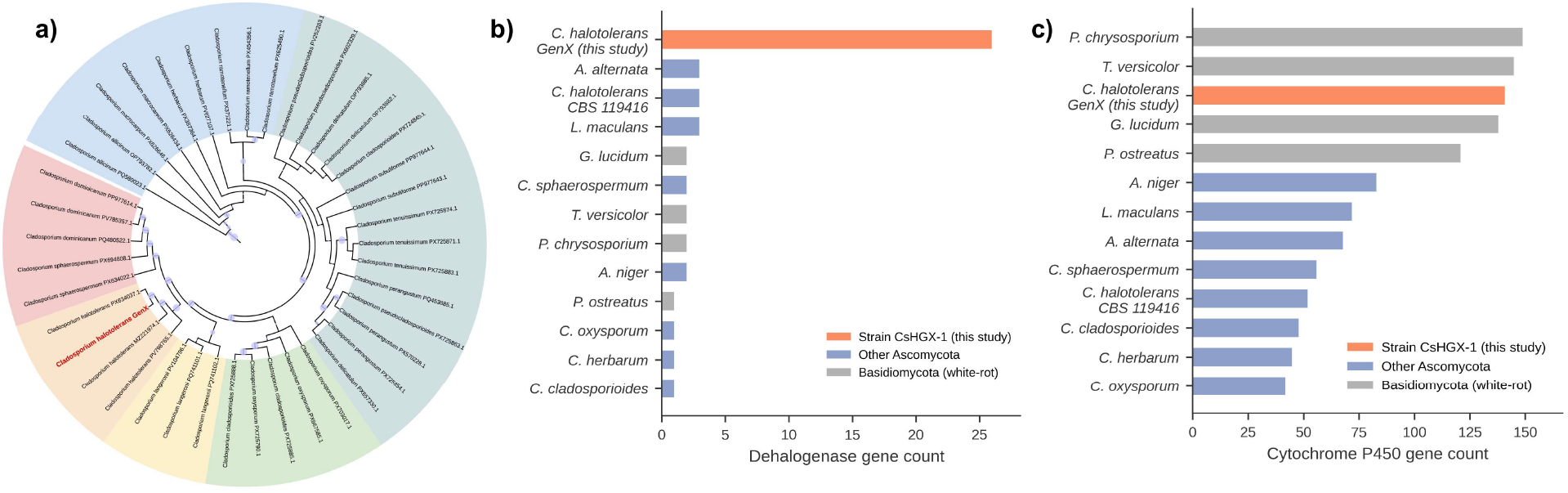
Comparative genomic enrichment of PFAS-degradation gene families. (a) Maximum-likelihood phylogenetic placement of strain CsHGX-1 (red) within the *Cladosporium halotolerans* species complex. (b) Dehalogenase and (c) cytochrome P450 gene counts across strain CsHGX-1 and 12 reference fungal genomes spanning Cladosporiaceae, other Ascomycota, and Basidiomycota white-rot taxa. Strain CsHGX-1 (orange) carries 8-fold more dehalogenases than its closest congeners.

### Aerobic GenX removal at 48 h is accompanied by measurable fluoride release

Cultures grown aerobically with GenX (50 mg L-1) as the sole carbon source removed 47.6 ± 1.7% of GenX (2,547.2 to 1,335.4 ng mL-1; LC-MS/MS) over 48 h, with no significant loss in abiotic controls (Figure 2a). Fluoride, measured by ion-selective electrode at the same time points, rose from baseline to 0.079 ± 0.018 mM exclusively in live cultures, confirming genuine C-F bond cleavage rather than physical or abiotic loss. For context, native microbial communities achieve under 6% HFPO-DA removal over 10 months under anoxic conditions, and the best-characterized anaerobic degrader, *Acidimicrobium* sp. strain A6, requires 120 days and external reductants to remove 12-57% of related PFAAs (11). A separate, longer resazurin-based metabolic assay, independent of the LC-MS/MS and RNA-seq time course and sampled daily over 5 days, showed metabolic activity in GenX-exposed cultures exceeding controls from day 1 onward (1.37-1.52-fold; p ≤ 0.003; Figure 2b), indicating sustained catabolic engagement rather than a transient stress response.

**FIGURE 2.**
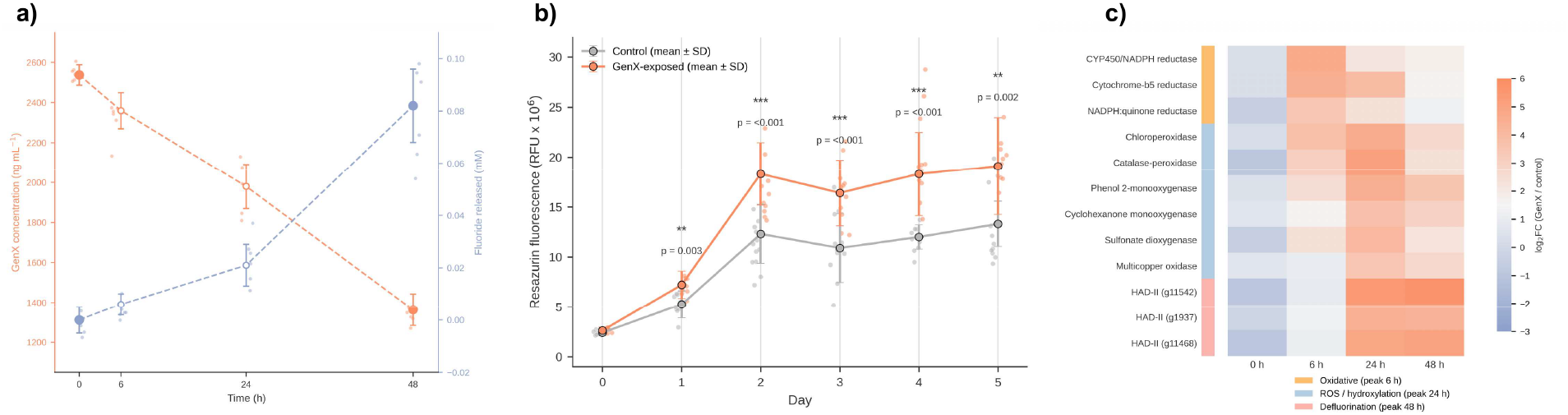
Transcriptional and metabolic response to GenX exposure. (a) GenX concentration (orange) and fluoride release (blue) over 48 h (mean ± SD). (b) Resazurin metabolic activity (mean ± SD relative fluorescence units) in GenX-exposed versus control cultures over 5 days, independent of the 48 h LC-MS/MS and RNA-seq time course. ** p < 0.01; *** p < 0.001. (c) Heatmap of log2FC (GenX/control) for the 12-gene PFAS-catabolism signature across four time points, grouped by phase of peak induction (oxidative, ROS/hydroxylation, defluorination).

### Time-resolved RNA-seq reveals a three-phase enzymatic program

RNA-seq across four time points (0, 6, 24, 48 h; n = 6 biological replicates per condition) resolved a tightly sequenced transcriptional response among 12 candidate PFAS-catabolism genes (Figure 2c; Table 1). At 6 h, oxidative genes, cytochrome P450/NADPH-P450 reductase (25.4-fold) and cytochrome-b5 reductase (22.7-fold), were induced first. At 24 h, induction shifted to hydroxylation and ROS-management genes, phenol 2-monooxygenase (22.8-fold) and catalase-peroxidase (33.5-fold; 15, 16). At 48 h, the three type-II HAD transcripts peaked (50.9-, 31.8-, and 20.9-fold), precisely when fluorinated intermediates generated by upstream oxidation would be expected to accumulate, coincident with measurable fluoride release (12, 13). This oxidation-first, HAD-mediated sequence is consistent with a model in which CYP450-driven hydroxylation of the ether linkage of GenX activates the molecule for subsequent hydrolytic defluorination (5, 14), and provides, to our knowledge, the first transcriptome-level evidence for this sequence in an Ascomycete fungus.

**TABLE 1.** Fold changes (GenX/control) of the 12-gene PFAS-catabolism signature across RNA-seq time points (n = 6 replicates per condition). Bold = peak time point per gene.

| Gene ID | Annotation | Phase (peak) | FC 0 h | FC 6 h | FC 24 h | FC 48 h |
| --- | --- | --- | --- | --- | --- | --- |
| g10139.t1 | CYP450/NADPH-P450 reductase | Oxidative (6 h) | **1.3** | 25.4 | 4.5 | 3.5 |
| g10866.t1 | Cytochrome-b5 reductase | Oxidative (6 h) | **1.2** | 22.7 | 15.4 | 3.1 |
| g602.t1 | NADPH:quinone reductase | Oxidative (6 h) | **0.7** | 10.9 | 5.0 | 2.5 |
| g4116.t1 | Chloroperoxidase | ROS/Hydrox. (24 h) | 1.2 | **13.9** | 24.2 | 6.4 |
| g1660.t1 | Catalase-peroxidase | ROS/Hydrox. (24 h) | 0.5 | **8.4** | 33.5 | 5.0 |
| g10740.t1 | Phenol 2-monooxygenase | ROS/Hydrox. (24 h) | 1.4 | **5.7** | 22.8 | 11.8 |
| g7946.t1 | Cyclohexanone monooxygenase | ROS/Hydrox. (24 h) | 1.6 | **3.0** | 13.2 | 8.0 |
| g5227.t1 | Sulfonate dioxygenase | ROS/Hydrox. (24 h) | 0.7 | **5.0** | 17.4 | 4.8 |
| g11976.t1 | Multicopper oxidase | Auxiliary | 1.6 | 2.0 | 11.2 | 6.7 |
| g11542.t1 | 2-Haloacid dehalogenase (HAD-II) | Defluorination (48 h) | 0.5 | 1.6 | **41.5** | 50.9 |
| g1937.t1 | 2-Haloacid dehalogenase (HAD-II) | Defluorination (48 h) | 0.9 | 2.5 | **20.9** | 20.1 |
| g11468.t1 | 2-Haloacid dehalogenase (HAD-II) | Defluorination (48 h) | 0.5 | 2.2 | **29.3** | 31.8 |

HAD-II, type-II haloacid dehalogenase. FC values calculated from raw counts (Supporting Information Table S1); full statistics in Table S3.

### Implications

Recovery of strain CsHGX-1 from activated sludge, an environment under continuous PFAS exposure, suggests that aerobic fungal defluorinators with expanded HAD repertoires may already be present, if overlooked, in engineered water systems (17, 18, 19). Its ability to use GenX as a sole carbon source under oxic conditions, without anaerobic cofactors, distinguishes it from previously characterized microbial PFAS degraders and makes it directly compatible with conventional aerobic treatment infrastructure. In 2024, the U.S. EPA added GenX to its list of RCRA hazardous constituents, underscoring the regulatory relevance of this finding (20). Resolving the identity of fluorinated intermediates and establishing a complete fluorine mass balance across the full-time course are priorities for confirming the complete defluorination pathway.

## MATERIALS AND METHODS

### Strain isolation and identification

*C. halotolerans* strain CsHGX-1 was isolated from activated sludge (Texas A&M University wastewater treatment plant, College Station, TX) by selective enrichment on M9 minimal salts medium containing GenX (100 mg L-1) as the sole carbon source. Taxonomic identity was confirmed by internal transcribed spacer (ITS) rDNA sequencing (primers ITS1/ITS4) and maximum-likelihood phylogenetic placement (1,000 bootstrap replicates; 41 accessions spanning 15 *Cladosporium* species).

### Genome sequencing and annotation

Genomic DNA was sequenced by Plasmidsaurus (Eugene, OR) on Oxford Nanopore R10.4.1 flow cells (v14 chemistry; Dorado v5.0 high-accuracy basecalling), quality-filtered (nanoq v0.10.0), and assembled with Hifiasm v0.25.0. Genes were predicted and annotated with funannotate2, integrating ab initio predictors with protein/transcript evidence and functional databases (Pfam, dbCAN, MEROPS, SwissProt); assembly completeness was assessed with BUSCO v5.7.1. PFAS-relevant gene families were curated by HMM search against manually compiled dehalogenase and halogenase profiles.

### GenX and fluoride quantification

Cultures (M9 + 50 mg L-1 GenX, 25°C, 100 rpm, n = 6 biological replicates, with abiotic autoclaved controls) were sampled at 0, 6, 24, and 48 h. GenX was quantified by LC-MS/MS (Vanquish-Altis, Thermo Scientific; negative ESI; SRM 285.0 → 169.0/119.0; method quantification limit 1.5 ng mL-1; matrix-matched calibration, 1-5,000 ng mL-1, quality-control acceptance criteria ±15% of nominal concentration every 10 injections). Fluoride was measured by ion-selective electrode (PerfectION, Mettler Toledo) using the standard addition method (method detection limit 0.02 mg L-1). Complete chromatographic, calibration, extraction, and analytical validation parameters are provided in Supplemental Methods.

### Metabolic activity

A separate, longer-timescale resazurin fluorescence reduction assay (96-well format) was sampled daily over 5 days in parallel GenX-exposed and control cultures (10-12 outlier-filtered wells per group per day; two-sample t-test after IQR-based outlier removal).

### Transcriptomics

RNA-seq libraries (3′-end counting, poly(dT)VN priming with UMIs) were prepared and sequenced by Plasmidsaurus from the same 0/6/24/48 h cultures (n = 6 per condition). Reads were filtered (FastP), aligned (STAR v2.7.11), deduplicated (UMICollapse), and quantified (featureCounts) against the CsHGX-1 genome; differential expression used edgeR v4.0.16. Fold changes are reported as GenX mean counts/control mean counts for 12 candidate PFAS-catabolism genes. Full bioinformatic parameters and statistical tests are provided in Supplemental Methods; raw count matrices are in Table S1.

## SUPPORTING INFORMATION

Supplemental Methods (complete analytical, chromatographic, and bioinformatic parameters, including quality-assurance and quality-control criteria); Table S1, raw RNA-seq count matrix; Table S2, mean ± SE counts per condition and time point; Table S3, fold-change and p-values by gene and time point; Table S4, resazurin metabolic fluorescence kinetics (day 0-5); Table S5, comparative genomic distribution across 13 fungal genomes.

## ACKNOWLEDGMENTS

We thank the Departments of Plant Pathology and Microbiology, and of Biochemistry and Biophysics, at Texas A&M University for laboratory support. This study was supported by the National Institute of Standards and Technology (NIST) project “Bioenvironmental Security and Training Program” (award no. 60NANB24D118). We acknowledge Plasmidsaurus for genome and transcriptome sequencing services. The authors declare no competing interests.

## DATA AVAILABILITY

The genome assembly and annotation of strain CsHGX-1, raw RNA-seq count matrices, GenX/fluoride quantification data, resazurin metabolic kinetics, and comparative genomic data are deposited at Figshare (https://doi.org/10.6084/m9.figshare.33098837).

